# Spillover of managed bumble bees from Mediterranean orchards during mass flowering causes minor short-term ecological impacts

**DOI:** 10.1101/2024.09.07.611777

**Authors:** Nitsan Nachtom Catalan, Tamar Keasar, Chen Keasar, Moshe Nagari

## Abstract

Commercial bumble bee colonies are routinely used for crop pollination in greenhouses, and are increasingly introduced into orchards as well. Bumble bee spillover to natural habitats neighboring the orchards may interfere with local wild bees and impact the pollination of non-crop plants. Concurrently, foraging in natural habitats may diversify the bumble bees’ diets and improve colony development. To evaluate these potential effects, we placed commercial *Bombus terrestris* colonies in blooming Rosaceae orchards, 25-125 m away from the margins. We recorded the colonies’ mass gain, population sizes, composition of stored pollen, and temperature regulation. We monitored bee activity, and seed sets of the non-crop plant *Eruca sativa*, along transects in a semi-natural shrubland up to 100 m away from the orchards, with managed bumble bees either present or absent. Rosaceae pollen comprised ∼1/3 of the colonies’ pollen stores at all distances from the orchard margins. Colonies placed closest to the margins showed prolonged development, produced fewer reproductive individuals, and had poorer thermoregulation than colonies closer to the orchards’ center. Possibly, abiotic stressors inhibited the bumble bees’ development near orchard borders. Wild bees were as active during the colonies’ deployment as after their removal. *E. sativa*’s seed sets decreased after bumble bee removal, but similar declines also occurred near a control orchard without managed bumble bees. Altogether, we found no short-term spillover effects of managed bumble bees on nearby plant-bee communities during the orchards’ two-week flowering. The colonies’ prompt removal after blooming can reduce longer-term ecological risks associated with managed bumble bees.

## INTRODUCTION

Agricultural crop pollination depends considerably on commercial pollinators, primarily honey bee and bumble bee colonies that are temporarily introduced into agroecosystems during crop flowering periods (Geslin et al. 2017). Bumble bees, the focus of the present work, have been mass-reared for decades, initially mainly to pollinate greenhouse crops (Velthuis and van Doorn 2006). More recently, commercial bumble bee hives are also increasingly used to pollinate open-grown crops, such as Rosaceae orchards that bloom in early spring under unstable weather conditions. Originating from temperate-climate areas, bumble bees are effective thermoregulators that can maintain foraging at lower temperatures than other bees (Corbet et al. 1993; Dag et al. 2006). Adding commercial bumble bee hives to honey bee hives in apple, pear and cherry orchards substantially increases cross-pollination, fruit set, final yield, and fruit size (Dag et al. 2006; Sapir et al. 2017; Earaets et al. 2020).

Beyond the demonstrated benefits of managed bumble bees for agricultural yields, their activity can also spill over to nearby non-crop areas (Trillo et al. 2019, 2020). Spillover can affect the pollination service that the managed bees provide to the agricultural crops (Carturan et al. 2023). In addition, spillovers were proposed to influence the development of the managed bee colonies, as well as the functioning of natural pollination interactions. We review each of these potential effects in the next paragraphs.

### Spillover, diet diversification, and colony development

Balanced diet is critically important to bee health (Wright et al. 2018 for honey bees). Nutrient deficiency in bumble bee hives may cause them to forage for nutritionally complementary foods0. As a result, hives placed in monoculture orchards, near wild natural areas, may benefit from foraging on diverse pollen sources outside the orchards. Evidence for this hypothesis is equivocal. Bumble bee colonies that were placed in flower-diverse suburban habitats collected more diverse pollen, grew better and gained more weight than colonies placed in monoculture crop areas (Goulson et al. 2002). Similarly, pollen collection and colony growth of orchard-pollinating bumble bees improved with the availability of natural habitat around the orchards (Proesmans et al. 2019). The latter study, however, found no correlation between the composition or diversity of pollen collected by the bees and colony growth rates. Moreover, a controlled laboratory feeding experiment found no advantage to pollen mixtures over some single-species high-quality pollen in supporting *B. terrestris* colony development (Moerman et al. 2017).

### Spillover effects on wild pollinators

Wild pollinators are essential for the reproduction of the natural flora, and complement honey bee pollination in many crops (Garibaldi et al. 2013). Pollinator diversity conservation is therefore important for natural ecosystems as well as for crop production. Worldwide reports indicate declines in the diversity of wild pollinators, resulting from losses of nesting and foraging habitats due to urbanization and agricultural expansion; declines in the quality and diversity of the bees’ food sources; and increased usage of pesticides in agriculture (Dicks et al. 2021)0. Managed pollinators are another stressor of wild bees, because of exploitation competition for food sources and transmission of pathogens and parasites (Mallinger et al. 2017; Iwasaki and Hogendoorn 2022). A particular concern regarding some managed bumble bees, such as *B. terrestris*, is their potential to establish colonies outside of croplands (Geslin et al. 2017), become invasive, and hybridize with local populations (Bartomeus et al. 2020, but see Franchini et al. 2023). In some of the documented bumble bee invasions, extensive influences on the local plant and pollinator communities were observed (Dafni and Shmida 1996; Morales et al. 2013, Acosta et al. 2016). Bumble bees are considered dominant competitors of wild bees owing to their social life style, generalist diet and cold hardiness that allows them to forage on cold days and in the early morning hours (Corbet et al. 1993; Bloch et al. 2017). Most studies to date documented competitive interactions between the managed bumble bee *B. terrestris* and local bumble bee species (Iwasaki and Hogendoorn 2022), or even with local populations of *B. terrestris* (Trillo et al. 2019). Little is known about the impacts of managed bumble bees on other bee species, such as the solitary bees that dominate Mediterranean ecosystems.

Local *B. terrestris* populations in Israel were historically limited to high mountains in the north of the country. The species extended its distribution range in recent decades (Bar Shai et al. 2013). *B. terrestris*’ arrival to a new area (the Carmel Mountain) was accompanied by reduced activity of other bee species0. Other field observations confirmed that *B. terrestris* foragers visit many wild plant species over a long activity season in natural sites, starting at earlier morning hours than the local solitary bees (Bar Shai et al. 2013). These observations suggest a competitive potential between bumble bees and local bees over food sources. Nevertheless, the effects of bumble bees on the local bee fauna at the interface between agricultural and natural habitats have not yet been assessed. Detailed studies of *B. terrestris*’ current distribution in Israel are also still lacking.

### Spillover effects on non-crop plants

Spillover of managed bumble bees from crops to nearby natural areas can have complex effects on the pollination prospects of wild plants. Managed bees may provide complementary pollination services that non-crop plants require, and thereby enhance the reproduction of these plants. Managed pollinators indeed visit some wild plants at the margins of fields, and this increases these plants’ seed sets (Geslin et al., 2017). However, managed pollinators sometimes also disrupt the pollination of native plants by displacing their specialized local pollinators. For example, invasive *B. terrestris* reduced the seed set of native plants (*Corydalis ambigua* and *Vicia nigricans*) in Japan and Argentina, respectively, through nectar robbing and by outcompeting the plants’ co-evolved native pollinators (Dohzono et al. 2008; Chalcoff et al. 2022).

Here, we addressed these potential implications of bumble bee spillover in a Mediterranean agroecosystem. We examined the influence of commercial *B. terrestris* hives in cherry and apple (Rosaceae) orchards on wild bee activity, as well as on the seed set of a non-crop plant species, in semi-natural areas near the orchards. We also examined whether the hives’ distance from the orchards’ borders (which we expected to associate with spillover levels), correlates with the fraction of non-Rosaceae pollen collected by the bees and with indicators of colony development. Much research has focused on spillover effects of wild pollinators from natural to agricultural habitats, their contribution to crop pollination (Garibaldi et al. 2013), and how it can be increased (e.g., Gilpin et al. 2022 for cherry). Our study asks a complementary question, focusing on spillover effects of managed bees from the agricultural to the natural habitat and their effects on wild pollinators and plants.

## MATERIALS AND METHODS

### Study site

Four early-blooming sweet cherry and apple orchards in the north of Israel (33°18’01’’ N, 35 °76’95’’E) were selected in each of the study years, 2021 and 2022. The study site, located at 1050 masl, is characterized by cool Mediterranean climate, 930 mm annual rainfall, and volcanic soil. We included both cherries and apples in the experiment because the two crops flower at the same time and provide similar food reward to foraging bees, in terms of protein, lipid, and amino acid composition of their pollen (Vaudo et al. 2020; Jeannerod et al. 2022). The blooming time of the selected cultivars is the earliest among the crops with assisted bumble bee pollination in the study area.

The orchards were 0.7-2.0 ha in area. To minimize soil and weather variation, all the orchards were located up to 1 km distance from each other. The minimal distance between orchards was 300 m, less than *B. terrestris*’ estimated maximal flight range of 2.5 km (Hagen et al. 2011). Thus, we cannot rule out flights of managed bumble bees between the orchards and neighboring semi-natural plots. All orchards were managed by the same grower using similar agricultural practices. Three orchards were chosen, each year, as treatment plots that received commercial bumble bee and honey bee hives. We used an apple orchard as a control, with commercial honey bee hives but no introduction of bumble bees. The control apple orchard was situated ∼950 meters away from the experimental orchards’ block. In 2021, the treatment orchards contained cherries only. In 2022, the treatment orchards contained cherries (two orchards) or a cherry-apple mix (one orchard). All orchards contained flowering herbaceous plants between tree rows and along their edges, and bordered a semi-natural Mediterranean shrubland on one of their sides. The flowering non-crop plants were dominated by species from the families Fabaceae, Asteraceae, Brassicaceae and Ranunculaceae.

### Experiment layout

At each study site, experimental manipulations and observations were conducted along a line transect from the middle of the orchard and up to 100 m into the natural habitat (Figure 1). We placed the commercial bumble bee hives inside each treatment orchard, along the transect, at three distances from the margin inside the orchard (25, 75 and 125 meters in 2021; and 25, 50 and 100 meters in 2022). We refer to these locations as ‘border’, ‘intermediate’, and ‘middle’, respectively. We placed two casings containing three hives (a ‘triple-pack’) at each location. The total density of bumble bee hives was ∼8.5 ha^−1^, and the honey bee hive density was ∼3 ha^−1^ – the recommended and commonly used densities for these crops. We marked points at 0, 10, 50 and 100 m from the margin into the semi-natural habitat outside all experiment and control orchards. These points were used for bee activity sampling and for monitoring of plant reproductive success, as described below.

**Fig. 1:**
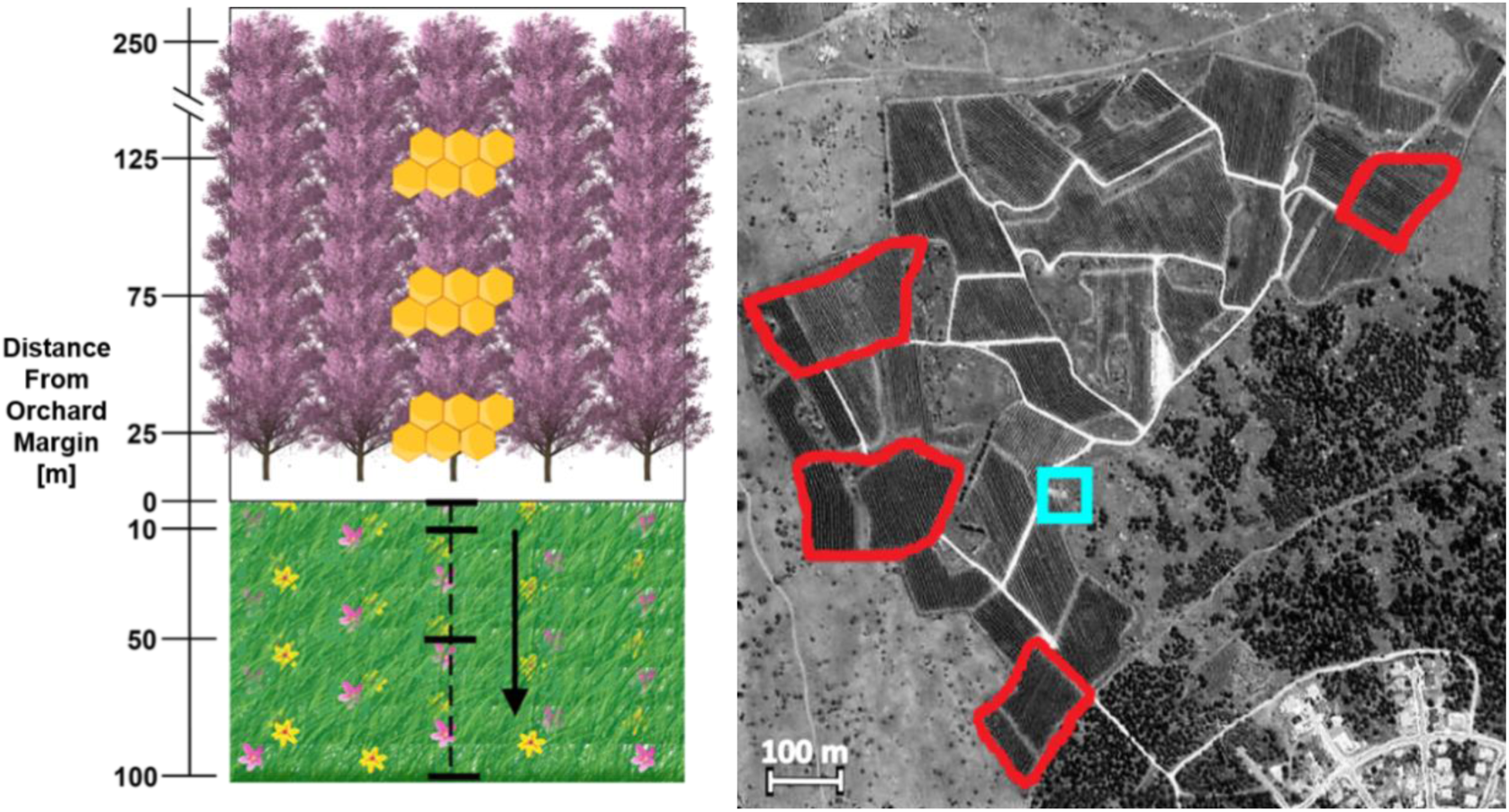
Study site – locations and experimental layout. An illustration of the experimental layout in one orchard in 2021. The orchard (top left panel) faces a semi-natural shrubland dominated by herbaceous plants (bottom left panel). Bumble bee hives (represented as hexagons) were placed at increasing distances from the orchard margin towards the orchard center at the beginning of apple and cherry blossom. The positions of pan traps and planters are indicated with thick horizontal lines. The dashed vertical line indicates the bee netting transect. The netting transect was divided into distance sections, also indicated with the horizontal lines. Right panel: Satellite photo of the study site, illustrating the treatment orchard plots in 2022 (red frames, left), the control orchard plot (red frame, top right) and the local meteorological station (light blue square).

### Colonies in the orchard

*B. terrestris* hives, each with a queen and about 100 workers and weighing 776.71 ± 3.04 g (mean ± SE, n=54 per year), were supplied by BioBee Ltd. (Sde Eliyahu, Israel). Upon arrival, we attached thermo-loggers (Maxim Integrated Products – iButton DS1921G-F5) to the bottom of 1-2 hives from each triple-pack. The loggers recorded the within-hive temperature once every 20 minutes, for 53 hives over the two study years. When placing the hives in the orchards, we made sure that their initial mass variation was minimal between orchards and distances from orchard borders (linear model: distance: F_2,75_ = 0.10, P = 0.91; orchard: F_5,75_ = 1.62, P = 0.16). A few days later, still before blooming, honey bee hives were placed throughout the orchards as part of the routine management practice for cherry and apple pollination.

Following hive placement, the bumble bees were allowed to exit and enter the hives freely until the end of the orchards’ bloom. On the last evening of the experiment, after all foragers returned to their colonies, we closed and weighed the hives, then stored them at −30°C until further processing.

### Colony demography and pollen stores

Bumble bees are primitively eusocial bees that live in annual colonies. Colonies are founded by a single fertilized queen, which rears the first batch of brood herself. When her daughters reach adulthood, they join her as sterile workers (Goulson 2003). The colony enters its eusocial phase and starts to grow rapidly. It first produces additional workers and then switches to the production of reproductive queens and drones, which eventually disperse and mate. The ratio between larvae and workers in the colony changes throughout the colony life in a predictable bimodal pattern (Duchateau and Velthuis 1988). Additionally, the number of reproductive individuals increases towards the end of the colony’s life. Thus, demographic data can provide information on colonies’ health, developmental stage, and reproductive success We counted the adult bees (classified into female workers, drones, and gynes) in 84 out of 108 of the frozen colonies. Immatures were classified into young brood cells (containing multiple eggs or young larvae in each wax cell up to their 3^rd^ instar), grown larvae (3^rd^ instar or older, which inhabit individual wax cells), small pupae (future workers or drones, ≤16 mm in length) and large pupae (future gynes, ≥ 18 mm in length). We also collected the stored pollen from each hive.

### Pollen composition analyses

#### Sample preparation

We analyzed the composition of stored pollen from a subsample of 34 colonies, placed in 2021 and 2022 in border (16 hives), intermediate (9 hives) and middle (9 hives) locations of two of the cherry orchards. We collected and mixed all the stored pollen from each colony and took 3-9 samples (depending on the amount of stored pollen) from the pollen mass for analysis. Each sample was diluted in water, spread on a microscope slide, dyed with methylene blue 1% and inspected under a light microscope at ×100 magnification. We photographed several (30-124) fields of vision from each slide, each containing at least 30 pollen grains. Overall, we obtained 1532 images containing ∼337,000 pollen grains.

#### Object detection

To identify and extract images of individual pollen grains, we used the faster-R-CNN (Ren et al., 2015) algorithm with pretrained parameters, downloaded from the detectron2 model-zoo (Wu et al., 2019). The faster-R-CNN algorithm learns from manually labeled examples. To generate these examples, we created a dataset of 15 images, in which we drew bounding boxes around all pollen grains, a total of 3184 grains. Given this dataset we applied the following protocol to generate and evaluate the object detection model:

1. The dataset is split to a train set (12 images, 2427 pollen grains) and a test set (3 images, 757 pollen grains).
2. The train set is augmented by rotating each image by 90°, 180°, and 270°, thus increasing the effective number of sampled pollen grains.
3. The train set images are split into tiles of 500X500 pixels. Neighboring tiles are 50% overlapping.
4. Edge grains, which are artificially cut, are removed by “painting” them with background color. Note that each of them also occurs as internal grains in neighboring tiles.
5. The tiles are fed to the detectron2 training process. We trained the model for 4 epochs. Longer training did not improve performance.
6. During inference, images are split to overlapping tiles, bounding boxes are predicted per tile, and overlapping bounding boxes of neighboring tiles are merged to provide the final set of detected pollen grains.

Detectron2 provides a confidence score to each of its predicted objects (pollen grains in this case). With a confidence threshold of 0.5, the algorithm was able to recover 94% of the pollen grains with 98% precision.

#### Classification

Aiming to estimate the fraction of cherry pollen grains within the total pollen mass of the bumble bee colonies, we re-trained a ResNet152 (He et al., 2016) model originally trained on the ImageNet dataset (Russakovsky et al., 2015). For the retraining we manually classified a training set of 480 images of cherry pollen grains, and 1782 images of non-cherry ones. Training converged after 10 epochs. The accuracy of the model was estimated to be 0.78 and 0.86 for cherry and non-cherry grains, respectively, using a test set of 221 cherry pollen grains and 822 non-cherry ones.

### Bee activity in the semi-natural habitat

We captured active bees in the semi-natural habitat at increasing distances from the orchard in each plot, while the commercial bumble bee hives were deployed in the experimental orchards and after their removal. In 2022, we also recorded whild bee activity before the introduction of the managed bumble bees. We netted bees in 2021 and 2022, and used pan traps in addition to netting in 2022.

For netting, we walked slowly along the 100 m transect outside each orchard and collected wild bees up to 1m on either side of the transect. Each transect walk lasted 30-60 minutes. Sighted bumble bees and honey bees were recorded but not captured. We recorded the distance range from the orchard (0-10m, 10-50m or 50-100m) at which each bee was sighted or netted. This protocol was repeated three times on each netting day (2021: 8:30, 11:00 and 15:00; 2022: 9:00, 12:00 and 15:00). Sampling times are reported in Table S1. To estimate the community composition of wild bees in the orchards’ vicinity, a subset of the netted bees (102 individuals caught near one orchard in 2021) were identified to the lowest possible taxonomic resolution.

For pan-trap sampling, we placed cream-colored plastic bowls at 0, 10, 50 and 100 m in the semi-natural area outside each orchard, along the sampling transect. Three water-filled traps were placed in each location, 1m apart. We set the traps in the morning (starting between 9:00 and 10:30) and at noon (starting between 12:00 and 13:30), and collected them three hours later. The pan traps were set up once a week in each orchard, during four weeks in 2022. This covered the periods before, during and after the deployment of the managed bumble bees. Since some traps were lost and some trapping sessions were disrupted, we ended up with an incomplete pan trap dataset, as detailed in Table S2.

### Seed production in the semi-natural habitat

In 2021 we used planted roquette (*Eruca sativa*: Brassicaceae) to evaluate seed production at increasing distances from the orchard. Roquette is exclusively insect pollinated, locally abundant and produces many pods and seeds gradually over time. Flowers develop along vertical inflorescences, from the bottom upwards. This allows distinguishing between flowers that bloomed while bumble bees were present and flowers that bloomed after hive removal. Planters with roquette in early flowering were placed in the semi-natural shrubland near the orchards, at 0, 10, 50 and 100 m from the orchard margin. One planter was placed at each distance in each of the treatment orchards, and two planters at every point in the control orchard, for increased control sample size. Once the commercial bumble bee hives were deployed, we removed all the blooming and wilted flowers from the roquette inflorescences, and left the netted planters without further observation for 20 days, until the end of the orchards’ bloom and removal of the managed bees. During this time, any new flowers could be pollinated by managed bumble bees, honey bees, and wild bees. We then removed all the receptive flowers, marked each branch at the top flower removal point with a colored ribbon, and allowed new flowers and pods to develop above the ribbon for 10-11 additional days. Potential pollinators at this stage included only unmanaged bees. Finally, we removed the planters from the field site, clipped the inflorescences’ heads to prevent additional blooming, and allowed the plants to age and dry naturally. We counted the flowers (flower stem remains) and seeds that developed while and after bumble bees were placed in the orchards.

### Data Analysis

We conducted all data analyses in R (R Core Team, 2022), using the packages ‘lme4’ (Bates et al., 2015), ‘MASS’ (Venables and Ripley, 2002), ‘plotrix’ (Lemon, 2006), and ‘stringr’ (Wickham, 2022).

#### Colony development

We used Generalized Linear (GLM) models to test the effects of hive location (border, intermediate or middle) in the orchards, year (2021 or 2022), and year × location interaction on the following variables: the total number of individuals, the numbers of egg/young larvae cells, larvae, small pupae, large pupae, adult workers, drones, gynes, total reproductives (drones + gynes + large pupae), the ratio of mature larvae to workers, and colony mass change (final mass - initial mass). The numbers of individuals in the different population groups, which are over-dispersed count data, were analyzed with negative binomial models. The larvae/worker ratio was analyzed using a Gamma family model with a log link function. Mass change data were analyzed with a Gaussian family model and an identity link function following a square-root transformation. We first ran the full models in all the analyses and then compared them to reduced models with fewer explanatory variables, using likelihood ratio tests. Different orchards within a single large block of Rosaceae plantations were included in the experiment in 2021 and in 2022. We are therefore unable to distinguish between the effect of year and of orchard identity in the statistical models.

#### Pollen composition

Each pollen grain in the dataset (n=337,162) was classified as ‘Rosaceae’ or ‘non-Rosaceae’. We used a Generalized Linear Mixed Model for binomially distributed data and a logit link function to test the effect of the hives’ distance from the orchard margins on the frequency of Rosaceae pollen. Hive identity was included as a random factor to account for the dependence between pollen grains that originate from the same hive.

#### Colony thermoregulation

We calculated the mean absolute difference between ambient and within-hive temperatures for each bumble bee colony, for the whole period of the colonies’ deployment in the orchard. Larger mean absolute differences indicate better ability of the bees to control the hive’s temperature. We used a General Linear Model for Gaussian distributed data and an identity link function to test the effects of distance from the margins, year, and their interaction on the mean temperature differences.

#### Bees in the semi-natural habitat

We used goodness of fit chi-square tests to assess whether wild bees were similarly active in different sections of the netting transect. The transect sections (0-10, 10-50 and 50-100 meters) differed in length. Therefore, if bees are equally active along the transect, the proportions of captures in the three sections are expected to be 0.1, 0.4 and 0.5, respectively. Pan traps were placed at 0, 10, 50 and 100 m from the orchard border. We compared the total numbers of bees caught at each of the four locations to a uniform expected distribution, namely 0.25 of all captures at each trapping location.

We also conducted chi-square tests for independence to examine whether the distribution of captured wild bees in the semi-natural habitat depended on the presence of managed bumble bees in the orchards. We compared the wild bees’ distribution in the experimental orchards between the periods of bumble bee presence and absence, separately for each year of the study and for each monitoring method. In 2022, we also compared the distribution of captured bees between the experimental orchards and the control orchard, for the period of bumble bee residence in the experimental orchards.

#### Seed production

To test the commercial bumble bees’ influence on pollination services for wild plants around the orchard margins, we used the ratio of seeds to flowers as a measure of pollination success. Using a GLM with the gamma family and a log link function, we tested the effects of orchard type (experiment/control), stage of the experiment (during/after bumble bee presence), and distance from the orchards’ margin on the number of seeds produced per flower.

## RESULTS

### *B. terrestris* colony development

The colonies’ mass change, total number of individuals, and numbers of individuals in most population subgroups (brood cells, small pupae, large pupae, workers, drones, and gynes) did not change consistently with distance from the orchard margins (Supplementary Table S3). The number of grown larvae (> 3^rd^ instar), on the other hand, declined from the margins to the center in both study years (Fig. 2, top panel, GLM: χ ^2^_2_ = 7.22, P = 0.03 for location, χ ^2^_1_ = 8.83, P = 0.003 for year, χ ^2^_2_ = 0.46, P = 0.80 for the interaction). Accordingly, the grown larvae to worker ratio was highest in the border hives in both years (Fig. 2, middle panel, χ^2^_2_ = 8.48, P = 0.01 for location, χ^2^_1_ = 41.63, P < 0.001 for year, χ^2^_2_ = 0.16, P = 0.92 for the interaction). Higher ratios indicate an earlier stage of the colony eusocial phase (Duchateau and Velthuis 1988). The number of reproductive individuals (gyne pupae + adult gynes + adult drones) was lower at the border of the orchard than in middle locations (Fig. 2, bottom panel), consistent with slower development of the colonies in border locations and/or lower reproductive success. The year, the distance from the orchards’ margin and the interaction between them significantly affected the number of reproductives (GLM: year: χ2 = 14.006, P-value = 0.0002; distance: χ2 = 4.285, P-value = 0.038; year and distance: 4.295, P-value = 0.038).

**Fig. 2:**
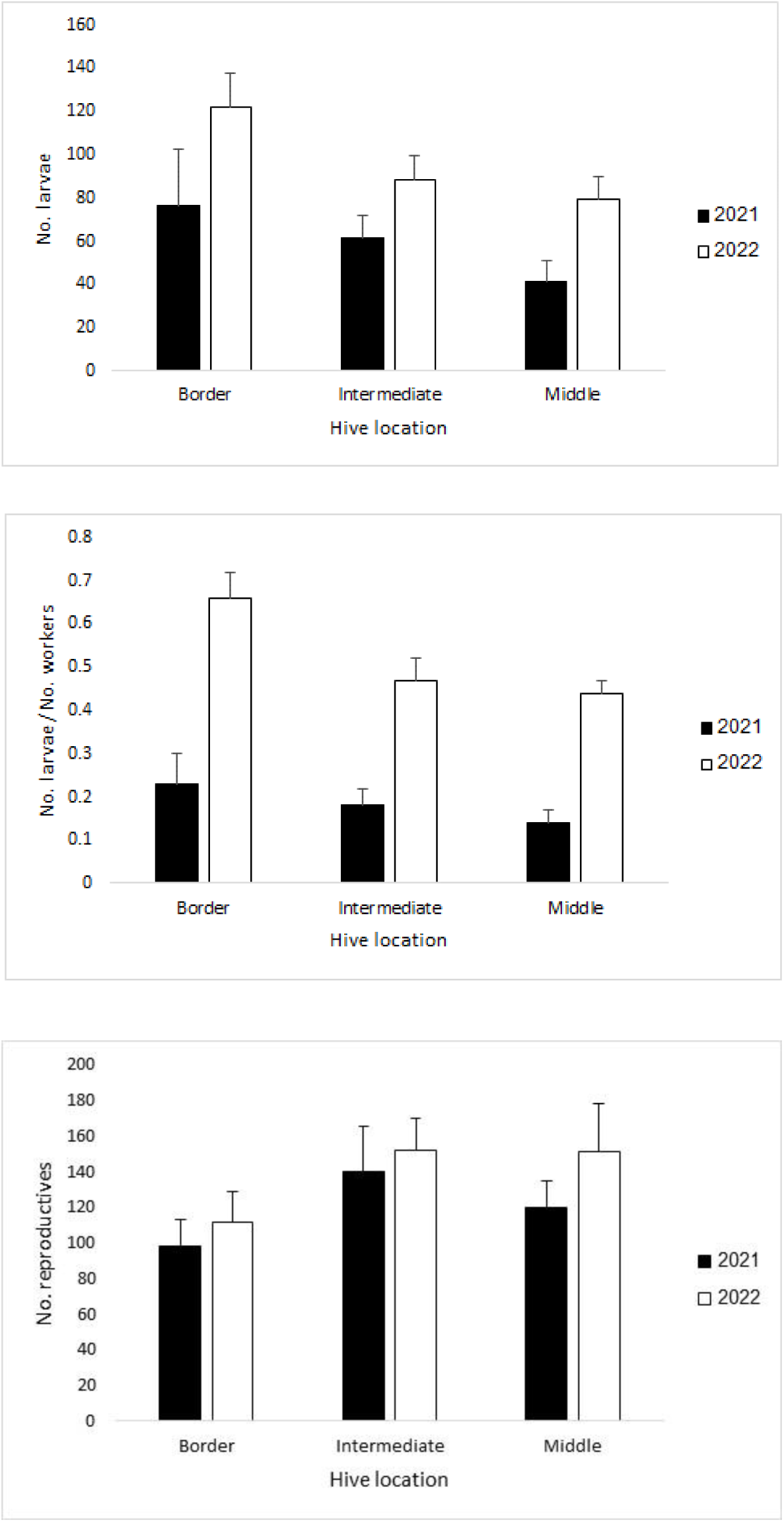
The effect of distance from orchard margins on colony demography. Bars indicate mean + SE of the number of mature larvae (top panel), larvae-to-worker ratios (middle panel), and the number of reproductive individuals (bottom panel) in the two study years. Border: orchard and shrubland border; Intermediate: inside the orchard, 50/75 m from the border; Middle: inside the orchard, 100/125 m from the border.

### Pollen composition

The mean±SE proportion of Rosaceae pollen in the samples of hive-stored pollen was 0.338±0.003 over all hives combined, 0.354±0.006 in the border colonies, 0.316±0.006 in the intermediate colonies, and 0.329±0.007 in the middle colonies. The distance from the margins did not significantly affect the proportion of stored Rosaceae pollen (logistic regression: χ^2^_5_ = 6.563, P-value = 0.255). These estimates are based on automated image classification and suggest that the Rosaceae fruit trees were not the major pollen sources for the bumble bees. Manual identification of a small random subsample of the data set (200 pollen grains) generated an even lower estimate of the proportion of Rosaceae pollen (19% of all grains).

### Colony thermoregulation

The mean±SD ambient temperature during the study was 15.2±7.1 C (range: 1.1-30.2 C), while in-hive temperatures were 27.1±2.6 C. The mean absolute environment-hive temperature difference tended to be lower in border locations than in the intermediate and middle locations, in both years (Fig. 3), suggesting less efficient thermoregulation at the orchards’ borders (GLM followed by likelihood ratio tests, effect of location: F_49, 51_ = 2.99, P = 0.06). Year and the location×year interaction did not significantly affect the colonies’ thermoregulation (year: F_47, 49_ = 0.52, P = 0.48, interaction: F_49, 50_ = 0.58, P = 0.56). Because thermoregulation is performed by the workers, we expected better temperature control in colonies with more workers. Indeed, the mean colony-environment temperature difference was positively correlated with the final number of workers (2021 and 2022 combined: N = 44, r = 0.43, P = 0.004, Supplementary Figure S4).

**Fig. 3:**
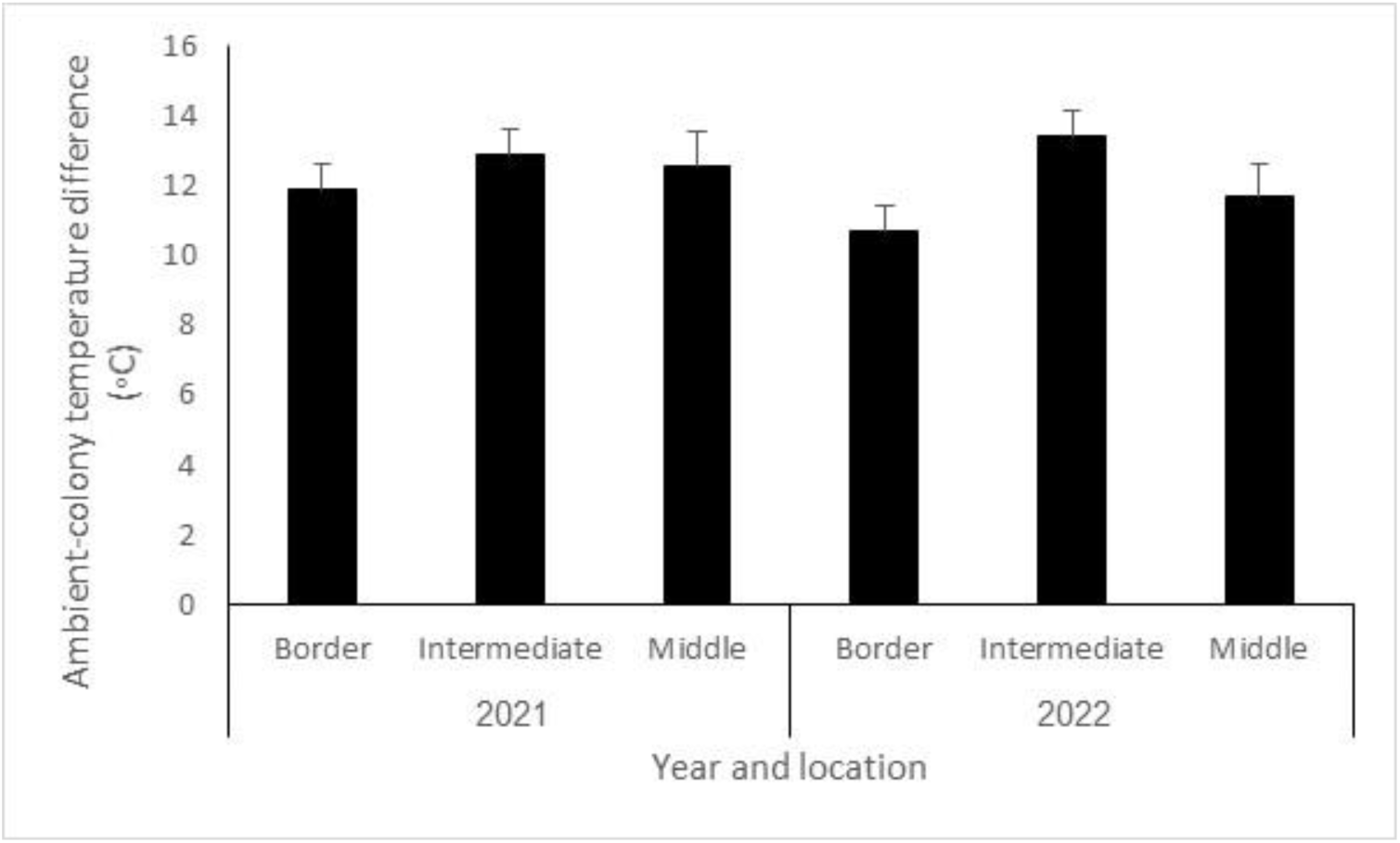
The effect of distance from orchard margins on colony thermoregulation. Bars indicate the mean + SE of the absolute temperature difference between the hives and the outside environment in the two study years. Border: orchard and shrubland border; Intermediate: inside the orchard, 50/75 m from the border; Middle: inside the orchard, 100/125 m from the border.

### Bee activity in the semi-natural habitat

The numbers of wild bees netted outside of the orchard fluctuated along the season, and so did the ambient temperature (Fig. 4). The identified subsample of 102 wild bees comprised 50.49% *Andrena*, 16.50% Osmini, 13.59% *Eucera*, 5.83% Dioxyni, and 13.59% individuals that we were not able to determine to family or tribe level. We recorded similar numbers of wild bees with or without the bumble bee colonies: there were 8.56±1.25 (mean±SE) wild bees per transect while the bumble bee colonies were inside the orchards, and 8.00±1.06 wild bees per transect after the bumble bees were removed. Of the identified wild bees, three *Andrena* species were dominant during the period of bumble bee deployment in the orchards. One species of the tribe Osmini dominated the assemblage after the bumble bees were removed. In 2021, we observed 0.44±0.11 bumble bee individuals per transect while the colonies were in the orchards and 0.58±0.25 individuals after the colonies were removed. We suspect that these individuals originated from other orchards or from nearby wild colonies. Honey bees and bumble bees were not documented in the transects in 2022, but bumble bees caught in the pan traps were counted. Four bumble bees were caught in the pan traps in total, three of them before the beginning of the orchards’ bloom and the deployment of commercial hives, and one after the end of bloom and hive removal. Honey bee activity along the transects in 2021 was much higher than bumble bee activity. We observed 10.11 ± 2.56 honey bees per transect walk when both honey bee and bumble bee hives were in the orchard, and 19.17 ± 3.19 honey bees after all the commercial hives were removed.

**Fig. 4:**
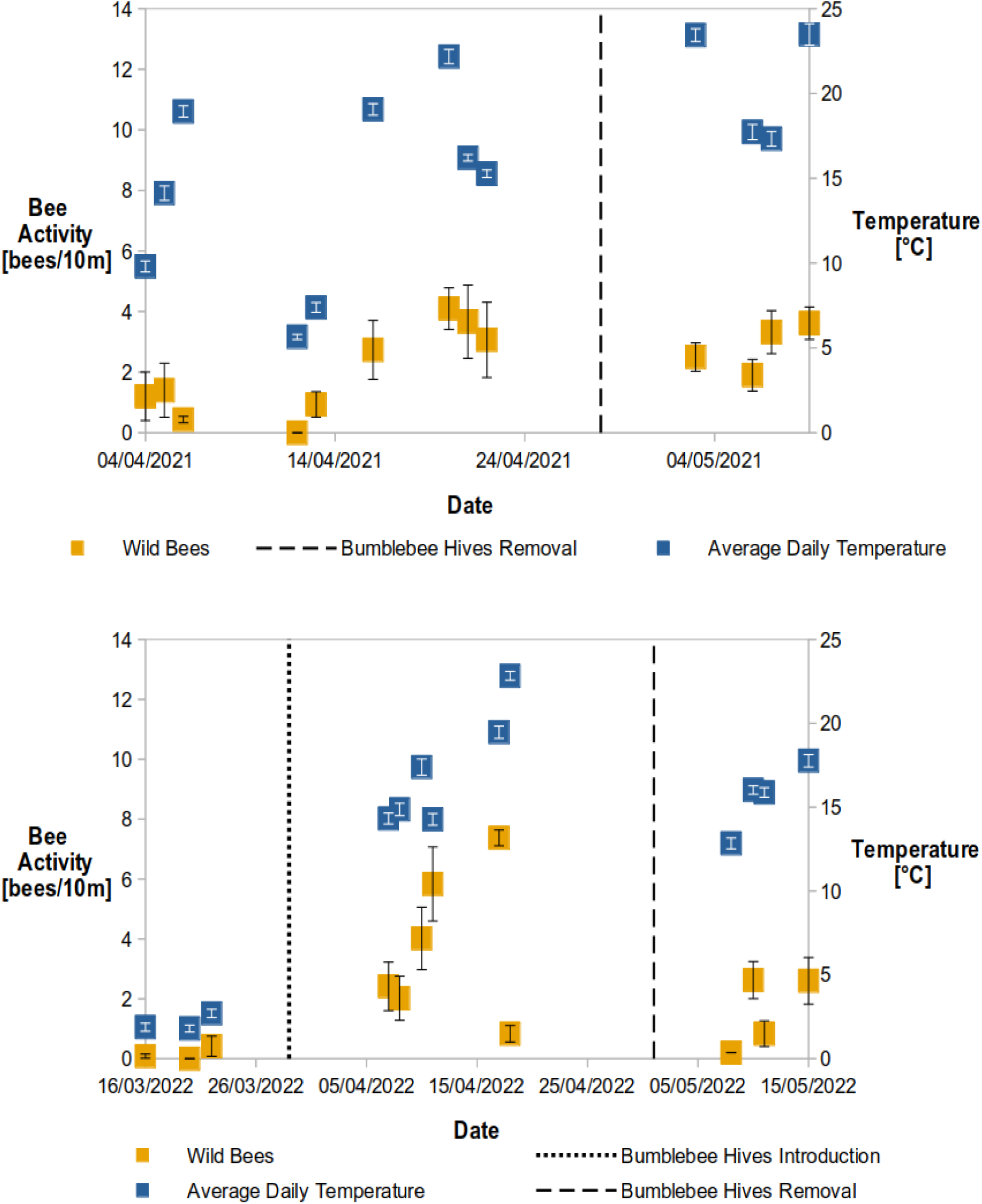
The relation between wild bee abundance outside the orchard and environmental temperature. top: 2021; bottom: 2022. Yellow markers indicate mean± SE numbers of wild bees caught per 10m transect. Blue markers indicate the daily (24 h) mean±SE ambient temperatures. Commercial bumble bee hive introduction (black dotted line) and removal (black dashed lines) dates are marked. In 2021 we did not sample wild bee activity before the introduction of bumble bee colonies.

Next, we asked whether wild bees were as active close to the orchards’ margins as further away from them. Altogether, we recorded 537 individuals in the transect walks over the two study years. These wild bees were captured near the treatment and the control orchards, both in the presence and in the absence of the managed bumble bees. 9.7% of these individuals were netted within 10 m of the orchard margins, 37.4% between 10-50 m, and 52.9% between 50-100 m from the margins. This distribution of captures does not differ significantly from the proportions of 10%, 40% and 50%, which are expected if wild bee activity in the natural area is unaffected by the distance to the orchard (Chi-square test for goodness of fit: χ^2^_2_ = 1.89, P=0.39).

Finally, we tested whether the distribution of wild bee captures along the netting transect depended on the presence of bumble bee colonies inside the orchards. Specifically, if bumble bees deter wild bees close to the orchard margins, we may expect a lower fraction of wild bee activity near the orchards while bumble bees were present than in their absence. To test this hypothesis, we compared the total numbers of wild bees trapped at each distance from the orchard borders, in the presence and absence of the managed bumble bees, separately for netting (Fig. 5 top) and for pan trap captures (Fig. 5 bottom). In 2021, the distribution of wild bees netted along the transects was independent of the monitoring period (with bumble bees in the orchards vs. after their removal, χ^2^_2_=1.54, P=0.46; Figure 5, top). In 2022, more of the wild bees’ netting captures were at 0-10 m of the orchards while bumble bees were present in the orchards than after their removal (χ^2^_2_=9.75, P=0.007). During the bumble bees’ deployment period, more wild bees were captured near orchards that contained bumble bees than to the control orchard (χ^2^_2_=11.36, P=0.003). These trends are opposite from the expectation of bumble bee interference with wild bee activity near the orchard margins. The distance distribution of wild bees captured in pan traps was independent of the presence of bumble bees in the orchards (Figure 5, bottom; experimental orchards during vs. after commercial bumble bee residence: χ^2^_3_=6.75, P=0.08; experimental orchards vs. control orchard during the bumble bees’ residence: χ^2^_3_=5.69, P=0.12). Taken together, we find no evidence for reduced activity of wild bees close to the orchards in the presence of managed bumble bees.

**Fig. 5:**
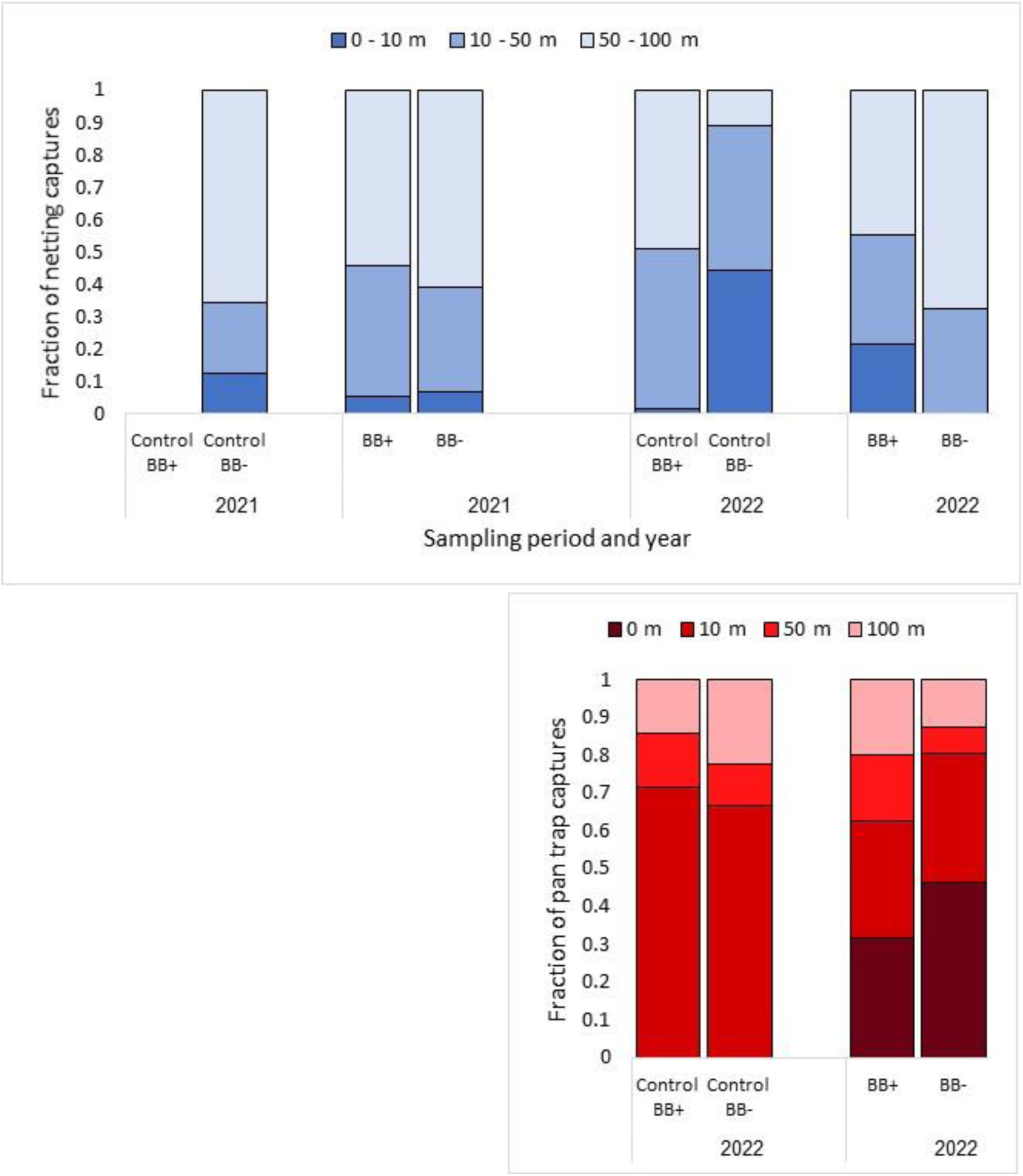
The effect of distance from the orchard and the presence of commercial bumble bee hives on wild bee abundance. Top, blue bars: the proportions of wild bees captured in the netting transects. Bottom, red bars: The proportions of wild bees captured in pan traps. Increasing distances are visualized as lighter shades. BB+ and BB-indicate the time periods in which commercial bumble bee hives were present in the experimental orchards and after their removal, respectively. The control orchards did not contain managed bumble bee hives throughout the whole study. In 2021, wild bees were netted near the control orchard only after the bumble bees’ residence period.

### Reproductive success of *E. sativa* plants

There were fewer seeds per flower after the bumble bees’ removal than in their presence, in the experimental orchards as well as in the control orchard (Fig. 6, GLM, effect of experimental stage: F_1, 251_=84.72, P<0.0001). This may indicate a seasonal decline in the plants’ reproductive success. Seed sets were higher in the control orchard than in the experimental orchards (treatment effect: F_1, 249_=5.02, P=0.03) in both stages of the experiment. Since there was no significant interaction between treatment and season (F_1, 247_=0.16, P=0.69), we cannot attribute the higher reproductive success in the control plot to the absence of bumble bees. Neither distance from the margin as a main effect nor any of the interactions significantly affected the variation in seed sets.

**Fig. 6:**
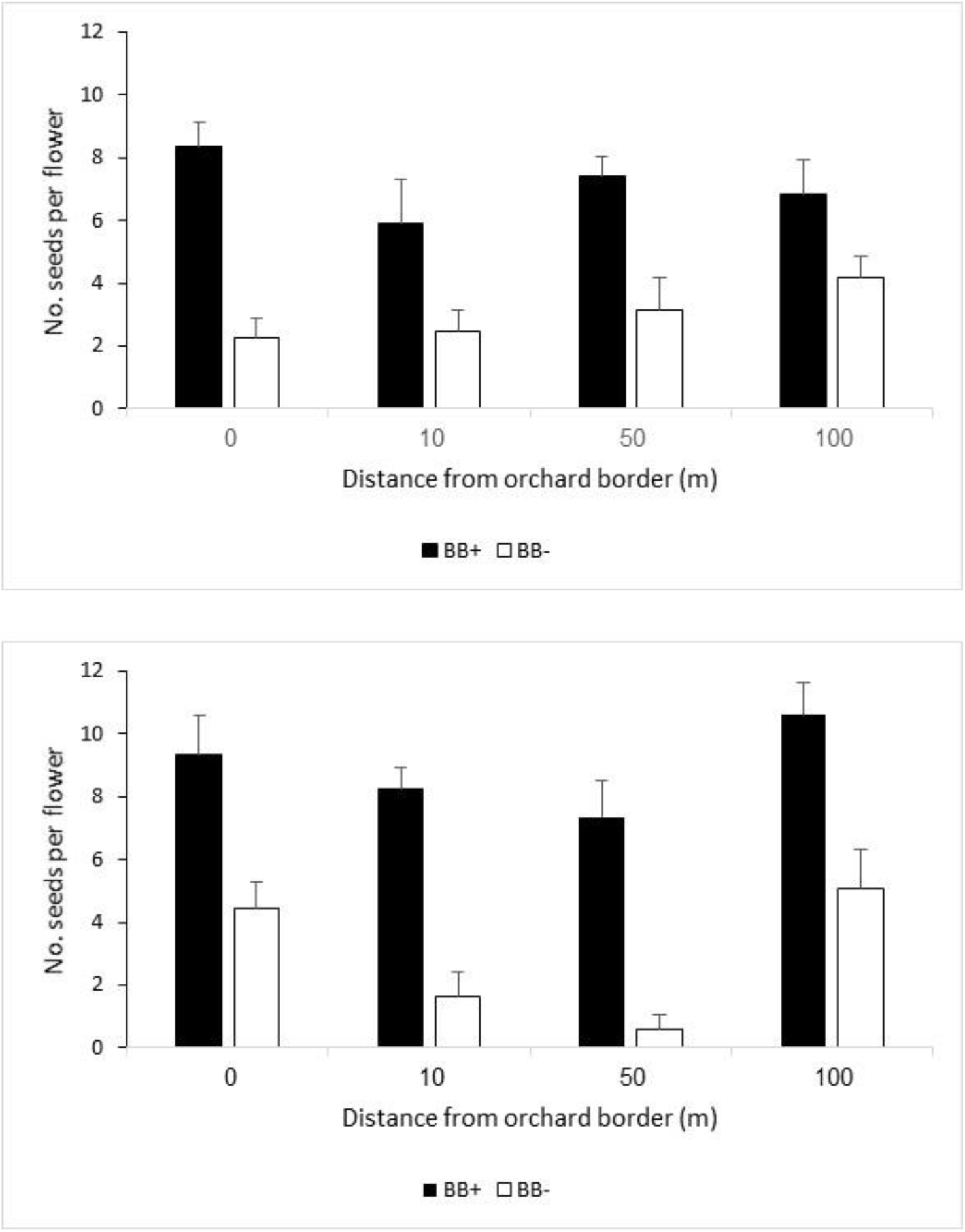
The effect of distance from the orchard and the presence of commercial bumble bee hives on *E. sativa* seed sets. Bars indicate the mean ± SE numbers of seeds per flower produced by *E. sativa* in the semi-natural habitat, at different distances from the orchard border. Top – experimental orchards where bumble bees were deployed. Bottom – the control orchard that received no bumble bees. BB+ and BB-denote the periods of bumble bee deployment and after their removal, respectively.

## DISCUSSION

We addressed some potential effects of spillover of managed bumble bees from orchards to nearby semi-natural habitats: improved colony development via diversification of the bees’ diet; interference with wild bees in the natural habitat; and impacts on the reproductive success of non-crop plants outside of the orchards. Below, we evaluate the evidence for spillover and for its resulting effects.

Transect walks and pan trap sampling indicated low activity of *B. terrestris* outside the orchards in our study. This is consistent with Fijen et al. (2022), who temporarily introduced managed honey bees into buckwheat fields to provide crop pollination. The honey bees foraged primarily within the fields and did not compete with wild bees in a nearby natural area. When observing bumble bees in the semi-natural habitat, we were not able to distinguish individuals of the experimental orchard colonies from native / feral individuals, nor from managed bumble bees introduced into other orchards by the farmers. Nevertheless, we recorded lower bumble bee activity in the natural habitat while the colonies were placed in the orchards than after their removal. If there had been a massive spillover of managed *B. terrestris* from the orchards to the semi-natural habitat, this should have actually reduced bumble bee activity after hive removal. *B. terrestris* activity in Israel starts in the spring, peaks in early summer, gradually declines and eventually stops in late summer (Bar Shai et al. 2013). Such seasonal increase in bumble bee activity over the spring is compatible with our findings. Interestingly, about 2/3 of the stored pollen in the hives did not originate from Rosacea, suggesting that the bumble bees fed on multiple plant species despite their low activity in the semi-natural habitat. This finding concurs with an earlier study, which found that even bumble bees placed within agricultural enclosures (tomato greenhouses) collected ∼30% of their pollen from non-crop plants (Whittington et al. 2004). We speculate that at least part of the bumble bees’ foraging resources were non-crop herbaceous plants within the orchards.

We expected colonies from border locations to develop better than mid-orchard colonies, owing to their proximity to diverse pollen sources in the natural habitat. Contrary to this prediction, middle, intermediate and border colonies stored similar percentages of cherry pollen, and did not differ in most of the developmental parameters. This may be explained by the small area of the orchards, relative to *B. terrestris*’ flight range (Hagen et al. 2011). As a result, diverse food sources may have been as accessible to middle colonies as to border colonies. Additionally, controlled feeding experiments found that Rosaceae pollen is a good dietary resource for bumble bees, even if offered as the only pollen source (Moerman et al. 2017).

Duchateau and Veltuis and (1988) showed that larvae/worker ratios in laboratory-reared *B. terrestris* colonies decline after the initiation of worker reproduction (the competition point), during the late phase of colony development. Since all of our colonies passed the competition point (had worker egg cells laid on top of each other), we assume that lower larvae/worker ratio reflects more advanced colony development. The higher larvae/worker ratios and the lower numbers of reproductives in the border hives indicates that they reached a less developed stage than the mid-orchard hives. Alternatively, the smaller number of reproductives may reflect lower reproductive success. Further, the border colonies’ inner temperature was more affected by the environment, indicating that they also had poorer thermoregulation than the middle colonies. Given that workers constantly incubate the brood and that tight temperature control is crucial for brood development (Heinrich, 1972, 1974; Vogt, 1986; Crall *et al*., 2018; Nagari *et al*., 2019), this may partly explain the differences in development rates and reproductive success. Although the number of workers in the colony strongly correlated with thermoregulation, worker numbers did not differ significantly between border and middle colonies. The decrease in thermoregulation capacity is therefore not due to insufficient worker population. We suspect that abiotic stressors, such as wind or low temperatures, are better buffered in mid-orchard locations than near the margins, allowing quicker development and higher reproductive success.

We captured wild bees at different distances from the orchard borders to assess whether their activity decreased near bumble bee colonies. We did not find consistent effects of the presence of managed bumble bees, nor of the distance from the colonies, on the numbers of captured wild bees. We did find seasonal changes in wild bee activity, which may be attributed to fluctuations in temperature, availability of non-crop flowers, or other environmental factors. Interestingly, in 2022 we also found opposite trends in bee distribution between the two monitoring methods. Netting capture at the margins of the experimental orchards decreased after the commercial hives were removed, while pan trap captures increased (Fig. 5). Different sampling methods are complementary and may sample different bee species (Wilson et al. 2008). Since netting targets bees in the vegetation, while pan traps capture primarily low-flying bees near the ground, this finding hints at a change in the bees’ community composition. Indeed, our sample of identified wild bees was dominated by different species during the bumble bees’ residence in the orchards *vs.* after their removal.

Concerns were raised in the literature about the use of *B. terrestris*, a species with invasive potential and history (Dafni and Shmida 1996; Morales et al. 2013; Acosta et al. 2016), for crop pollination. In the short term, managed bumble bees were feared to disrupt the activity of native wild bees while placed in the orchards. In the longer term, commercial bumble bees’ offspring can spread and establish feral populations that might compete with and transmit pathogens to wild bees (Mallinger et al. 2017). We found little evidence for short-term interference in our study system. However, we note that we did not test for longer-term effects on native wild bee activity, as we removed the hives from the site shortly after the end of bloom. Hives left in the orchard beyond the crop blooming season may pose higher risks to local bees, because they are more likely to forage in the semi-natural habitat and to initiate feral colonies. Until proven to be harmless, it is advisable to remove the commercial bumble bee colonies promptly after the orchards’ bloom to reduce their longer-term ecological risks.

We did not document an effect of distance from the orchard, alone and in interaction with the presence of bumble bees, on the seed set of *E. sativa*, a non-crop plant. The control plot data demonstrates a strong decline in the plant’s reproductive success in late spring, which is unrelated to the bumble bee colonies’ presence, and that may reflect resource limitation or climate stress experienced by the plants.

In conclusion, managed bumble bees did not massively spill over to the semi-natural habitat during the Rosaceae mass-flowering period, and subsequently also did not impact local bees and plants in the short term. The hives’ location within the orchards affected their development rate, but was unrelated to the composition of in-hive stored pollen. Current management recommendations, which call for removal of the managed bees immediately after the end of the crop bloom, should be carefully followed to minimize the bees’ spillover and potential longer-term adverse environmental effects.

## ACKNOWLEDGEMENTS

This study was funded by the Nekudat Hen Foundation for Sustainable Agriculture and by a Haifa University – Shamir Institute research cooperation grant. Ronen Shafir, Dor Farkash, Roni Levinson and Dana Amon assisted with field and lab work. The Botanical Garden at Oranim College helped in growing the potted plants. BioBee Sde Eliyahu LTD supplied the bumble bee hives for the experiment. Bees collected during the study are deposited at the Margolin House Collections, Oranim College.

## Author Contributions

NNC, MN, TK designed the study. NNC and MN performed experiments. All authors analyzed the data. TK wrote the paper. All authors read and approved the final manuscript.

## Ethics approval

This experiment involved insects. No ethics approval is required.

---

**Table S1:** Transect Netting Schedule and Completion.

| Year | Plot | Time of day | Before<br>Commercial<br>Bees | Week 1 | Week 2 | Week 3 | After<br>Commercial<br>Bees |
| --- | --- | --- | --- | --- | --- | --- | --- |
| 2021 | Control | Morning | X | X | X | X | Complete |
|  |  | Noon | X | X | X | X | Complete |
|  |  | Afternoon | X | X | X | X | Complete |
|  | A | Morning | X | Complete | Complete | Complete | Complete |
|  |  | Noon | X | Complete | Complete | Complete | Complete |
|  |  | Afternoon | X | Complete | Complete | Complete | Complete |
|  | B | Morning | X | Complete | Complete | Complete | Complete |
|  |  | Noon | X | Complete | Complete | Complete | Complete |
|  |  | Afternoon | X | Complete | Complete | Complete | Complete |
|  | C | Morning | X | Complete | Complete | Complete | Complete |
|  |  | Noon | X | Complete | Complete | Complete | Complete |
|  |  | Afternoon | X | Complete | Complete | Complete | Complete |
| 2022 | Control | Morning | X | *RS - Com-<br>plete | X | X | Complete |
|  |  | Noon | X | *RS - Com-<br>plete | X | X | Complete |
|  |  | Afternoon | X | *RS - Com-<br>plete | X | X | Complete |
|  | D | Morning | Complete | Complete | X | X | Complete |
|  |  | Noon | Complete | Complete | Complete | X | Complete |
|  |  | Afternoon | Complete | Complete | Complete | X | Complete |
|  | E | Morning | Complete | *RS - Com-<br>plete | X | X | Complete |
|  |  | Noon | Complete | *RS - Com-<br>plete | X | X | Complete |
|  |  | Afternoon | Complete | *RS - Com-<br>plete | X | X | Complete |
|  | F | Morning | Complete | Complete | Complete | X | Complete |
|  |  | Noon | Complete | Complete | Complete | X | Complete |
|  |  | Afternoon | Complete | Complete | Complete | X | Complete |

\*RS - Rescheduled

**Table 2:** Pan Traps Schedule and Completion.

**Table Error! No text of specified style in document.2: Pan Traps Schedule and Completion**
| Plot | Time of day | Before Commercial Bees | Week 1 | Week 2 | After Commercial Bees |
| --- | --- | --- | --- | --- | --- |
| Control (2022) | Morning | X | 0 & 10 – done | X | Complete |
|  | Noon | X | 0, 10 & 100 – done | Complete | Complete |
| D | Morning | Complete | Complete | X | Complete |
|  | Noon | Complete | Complete | Complete | Complete |
| E | Morning | Complete | Complete | X | Complete |
|  | Noon | Complete | Complete | Complete | Complete |
| F | Morning | Complete | Complete | Complete | Complete |
|  | Noon | Complete | Complete | Complete | Complete |

**Table 3:** Colony demography.

| Location | Year | Total population | No. workers | No drones | No gynes | No. egg cells | No. larvae | No. small pupae | No. large pupae | Pollen mass (g) | No. reproductives | No. Larvae/ No. wd | Mass change |
| --- | --- | --- | --- | --- | --- | --- | --- | --- | --- | --- | --- | --- | --- |
| Border | 2021 | 859.53 ± 73.89 | 323.2 ± 22.1 | 23.4 ± 6.42 | 32.73 ± 6.28 | 178 ± 31.19 | 76.33 ± 25.94 | 183.6 ± 37.03 | 42.27 ± 12.36 | 58.11 ± 8.32 | 98.4 ± 14.94 | 0.23 ± 0.07 | 366.8 ± 57.29 |
| Intermediate | 2021 | 821.83 ± 82.16 | 382.33 ± 57.08 | 21.83 ± 7.28 | 38.5 ± 11.4 | 119 ± 30.51 | 61.50 ± 10.64 | 119.17 ± 19.45 | 79.5 ± 17.29 | 67.12 ± 11.67 | 139.83 ± 25.66 | 0.18 ± 0.04 | 447.33 ± 88.37 |
| Middle | 2021 | 696.33 ± 94.24 | 297.11 ± 49.75 | 47.67 ± 12.05 | 24.56 ± 5.53 | 63 ± 8.45 | 41.56 ± 9.76 | 175.11 ± 40.3 | 47.33 ± 13.26 | 45.71 ± 8.33 | 119.56 ± 15.13 | 0.14 ± 0.03 | 356.89 ± 64.99 |
| Border | 2022 | 737.18 ± 66.25 | 180.41 ± 13.87 | 62.41 ± 16.48 | 9.41 ± 2.12 | 68.12 ± 10.22 | 122.18 ± 15.55 | 249.53 ± 33.97 | 45.12 ± 11.29 | 31.27 ± 6.94 | 116.94 ± 17.81 | 0.66 ± 0.06 | 450.53 ± 75.06 |
| Intermediate | 2022 | 776.83 ± 60.07 | 188.94 ± 16.26 | 65.28 ± 10.84 | 25.06 ± 6.53 | 52.22 ± 6.88 | 88.56 ± 11.4 | 294.72 ± 37.35 | 62.06 ± 13.06 | 25.62 ± 8.76 | 152.39 ± 17.53 | 0.47 ± 0.05 | 580.61 ± 78.35 |
| Middle | 2022 | 705.21 ± 70.89 | 190.05 ± 21.24 | 60.68 ± 10.96 | 24.89 ± 6.27 | 59.84 ± 9.3 | 79.26 ± 10.84 | 231.63 ± 29.52 | 58.84 ± 20.11 | 28.4 ± 9.51 | 144.42 ± 25.25 | 0.44 ± 0.03 | 503.58 ± 80.68 |

**Figure S4:**
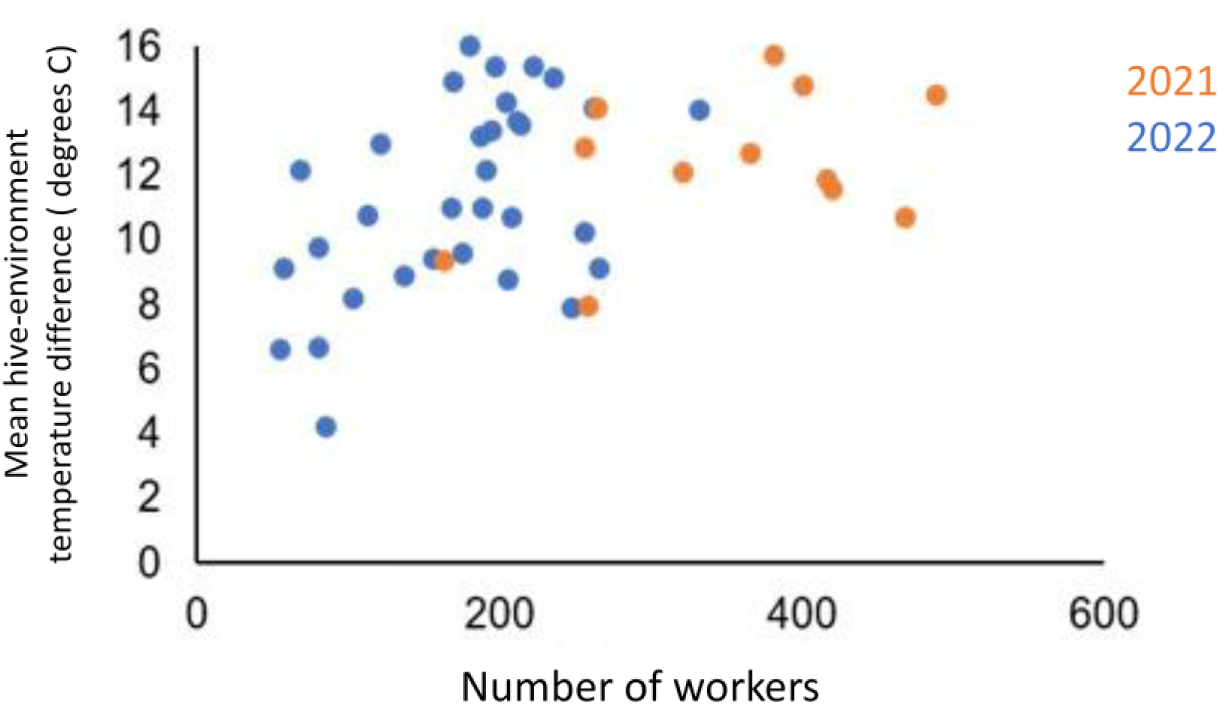
The mean temperature difference between the hive and the environment vs. the number of workers per hive.

